# Methylglyoxal-derived hydroimidazolone, MG-H1, increases food intake by altering tyramine signaling via the GATA transcription factor ELT-3 in *Caenorhabditis elegans*

**DOI:** 10.1101/2022.08.18.504374

**Authors:** Muniesh Muthaiyan Shanmugam, Jyotiska Chaudhuri, Durai Sellegounder, Amit Kumar Sahu, Sanjib Guha, Manish Chamoli, Brian Hodge, Charis Roberts, Gordon Lithgow, Richmond Sarpong, Pankaj Kapahi

**Affiliations:** The Buck Institute for Research on Aging, 8001 Redwood Boulevard, Novato, CA 94945, USA; Department of Chemistry, University of California, Berkeley, CA 94720, USA; Department of Urology, University of California, 400 Parnassus Avenue, San Francisco, CA 94143, USA

**Keywords:** Feeding, Advanced Glycation End-products (AGEs), *glod-4*, MG-H1, tyramine, GATA transcription factor, *elt-3*, *tyra-2*, *ser-2*, *C. elegans*, neuronal damage, pharyngeal pumping

## Abstract

The Maillard reaction, a chemical reaction between amino acids and sugars, is exploited to produce flavorful food almost everywhere, from the baking industry to our everyday life. However, the Maillard reaction also takes place in all cells, from prokaryotes to eukaryotes, leading to the formation of Advanced Glycation End-products (AGEs). AGEs are a heterogeneous group of compounds resulting from the irreversible reaction between biomolecules and α-dicarbonyls (α-DCs), including methylglyoxal (MGO), an unavoidable byproduct of anaerobic glycolysis and lipid peroxidation. We previously demonstrated that *Caenorhabditis elegans* mutants lacking the *glod-4* glyoxalase enzyme displayed enhanced accumulation of α-DCs, reduced lifespan, increased neuronal damage, and touch hypersensitivity. Here, we demonstrate that *glod-4* mutation increased food intake and identify that MGO-derived hydroimidazolone, MG-H1, is a mediator of the observed increase in food intake. RNA-seq analysis in *glod-4* knockdown worms identified upregulation of several neurotransmitters and feeding genes. Suppressor screening of the overfeeding phenotype identified the *tdc-1*-tyramine-*tyra-2/ser-2* signaling as an essential pathway mediating AGEs (MG-H1) induced feeding in *glod-4* mutants. We also identified the *elt-3* GATA transcription factor as an essential upstream factor for increased feeding upon accumulation of AGEs by partially regulating the expression of *tdc-1* and *tyra-2* genes. Further, the lack of either *tdc-1* or *tyra-2/ser-2* receptors suppresses the reduced lifespan and rescues neuronal damage observed in *glod-4* mutants. Thus, in *C. elegans*, we identified an *elt-3* regulated tyramine-dependent pathway mediating the toxic effects of MGO and associated AGEs. Understanding this signaling pathway is essential to modulate hedonistic overfeeding behavior observed in modern AGEs rich diets.

## INTRODUCTION

Processed modern diets enriched with Advanced Glycation End-products (AGEs), formed by Maillard reaction, are tempting to eat but at the same time deleterious for health [1] [2] [3]. In 1912, a French Chemist L.C. Maillard, reported a reaction between glucose and glycine upon heating resulting in the formation of brown pigments [4]. Later the covalent bonds formed between carbohydrates and proteins during heating in a non-enzymatic browning reaction was named the Maillard reaction [5] [6]. Glycation is a part of the Maillard reaction, or browning of food, during cooking which enhances the taste, color, and aroma of the food to make it more palatable [4] [7]. Maillard reaction has revolutionized the food industry by playing an important role in food chemistry [8]; however, this reaction also results in the formation of adverse AGEs as well as toxic byproducts and one of the well-studied toxic byproduct is acrylamide [9] [10] [11].

Apart from food, AGEs are endogenously produced in cells when either reducing glucose or α-dicarbonyl compounds (α-DCs) (such as glyoxal (GO), methylglyoxal (MGO), 3-deoxyglucosone (3DG), etc.) and non-enzymatically react with biomolecules. α-DCs are unavoidable byproducts of cellular metabolisms, such as glycolysis and lipid peroxidation. AGEs include GO derivatives such as carboxymethyl lysine (CML) and glyoxal lysine dimer (GOLD). AGEs derived from MGO include hydroimidazolone (MG-H1), carboxyethyl lysine (CEL), and methylglyoxal lysine dimer (MOLD), and 3DG derivatives include 3-deoxyglucosone-derived imidazolium cross-link (DOGDIC), Pyrraline, etc. [12] [13] [1] [14]. The glyoxalase system utilizes glyoxalases enzymes, Glo1 and Glo2, and reduced glutathione (GSH) to detoxify α-DCs stress, especially MGO to lactate, in cytosol and nucleus. Differential expression levels of glyoxalases are reported in various disease conditions such as diabetes, hypertension, neurodegenerative disorders, anxiety disorders, infertility, cancer, etc., suggesting their role in exacerbating their pathogenesis [15]. Glo1 has been linked with several behavioral phenotypes, such as anxiety, depression, autism, and pain, among other mental illnesses [16]. Also, we have previously demonstrated increased neuronal damage in the *C. elegans glod-4* glyoxalase mutant model, which is shown to accumulate high levels of α-DCs and AGEs [17]. AGEs accumulate in long-lived proteins, such as collagen and lens crystallins [18]; further, quantifying the glycated form of hemoglobin (HbA1c) is utilized as a biomarker in diabetes [19]. It is vital to notice that AGEs are strongly implicated in aging and associated diseases such as obesity, diabetes, neurodegeneration, inflammation, cardiomyopathy, nephropathy, and other diseases [20] [1] [21]. Especially, neurodegenerative diseases have demonstrated a strong correlation between AGEs levels and pathogenesis.

Overconsumption of food and excessive availability of cheap, highly processed foods lacking nutritional qualities contribute to the obesity pandemic. Obesity leads to other complications like diabetes, hypertension, cancers, cardiovascular, inflammatory, and neurodegenerative disorders, among other non-communicable chronic diseases [22] [23] [24] [25] [26] [27] [28] [29]. Although identifying genetic loci linked to obesity improves treatment options [30], exploring other signaling pathways modulating increased feeding behavior and obesity is essential. Here we report that loss of glyoxalase system increased feeding behavior in *C. elegans* mediated by accumulation of AGEs. We also identified the mechanism for the observed phenotype and found that MG-H1 acts via the *elt-3* GATA transcription factor to partially regulate the expression of *tdc-1*, an enzyme that biosynthesis tyramine neurotransmitter, and its receptor, *tyra-2* as well as *ser-2*, to mediate adverse effects of AGEs such as increased feeding, reduced lifespan, and neuronal damages. This study is the first to identify the signaling pathway mediated by specific AGEs molecules downstream of MGO (such as MG-H1) to enhance feeding and neurodegeneration. Our study emphasizes that AGEs accumulation is deleterious and enhances disease pathology in different conditions, including obesity and neurodegeneration. Hence, limiting AGEs accumulation is relevance to the global increase in obesity and other age-associated diseases.

## RESULTS

### AGEs increases food intake and food-seeking behavior in *C. elegans*

Our initial observations revealed that *glod-4* glyoxalases enzyme mutants exhibit a significantly enhanced pharyngeal pumping than wild-type N2 animals (Figures 1A). This increase in pharyngeal pumping was consistent from day 1 (young adult, post-65 hours of timed egg-laying) till day 3 of adulthood (Figure 1A). We performed a food clearance assay to validate whether increased pharyngeal pumping was also accompanied by enhanced food intake (Figure 1B + Figure Suppl. 1A) and found increased bacterial clearance after 72 hours in *glod-4* mutants. Further, treatment with serotonin resulted in increased bacterial clearance in both wild-type N2 worms and *glod-4* mutants (Figure Suppl. 1B+1C), suggesting that AGEs mediated increased pharyngeal pumping is independent of the serotonin signaling [31]. These preliminary observations lead to the hypothesis that enhanced feeding in *glod-4* mutant worms is either mediated by endogenous accumulation of α-DCs or AGEs characterized previously [17] [32] [33]. To this end, we found that MG-H1 and CEL as potential MGO-derived AGEs to cause increased feeding. Just feeding MGO was not sufficient to increase the pharyngeal pumping rate (Figure Suppl. 1D). Time course analysis in wild-type N2 worms treated with MG-H1 showed that 24 hrs of MG-H1(150 µM) treatment was enough to increase pharyngeal pumping significantly (Figure 1C). A significant increase in bacterial clearance was observed after 72 hours (Figure 1D). In addition, we also demonstrated that MG-H1 regulates pharyngeal pumping rate in a dose-dependent manner (Figure 1E). Since MG-H1 is the product of arginine modification by MGO, we used arginine as a negative control for our MG-H1 treatment. We did not observe a significant difference between worms treated with arginine versus water versus PBS (Figure Suppl. 1E). In addition to food consumption, *glod-4* mutant exhibited a significantly increased preference towards food source OP50-1 at day 1 and day 3 of adulthood compared to wild-type N2 worms (Figure 1F + Figure Suppl. 1F). Furthermore, we noticed that wild-type N2 worms preferred exogenous MG-H1 compared to MGO when provided with bacterial food source *E. coli* OP50-1. We did not observe this phenotype in the *glod-4* mutant background (Figure 1G), suggesting that MG-H1 and *glod-4* null mutation increases feeding by overlapping mechanism.

**Figure 1:**
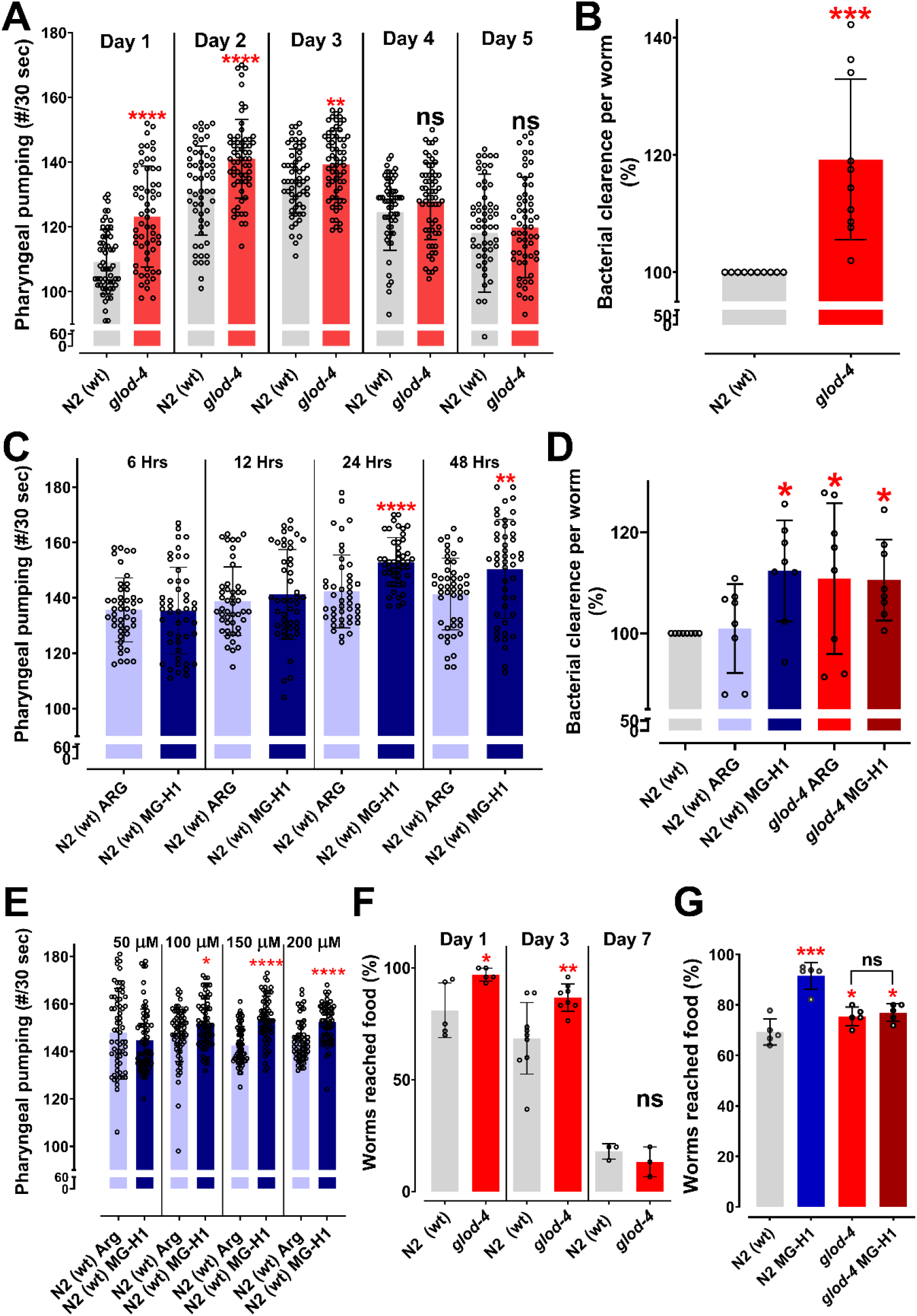
MG-H1 increases pharyngeal pumping and feeding in *C*. *elegans*. (A) Quantification of pharyngeal pumping (#/30 sec) in N2 (wt) and *glod-4 (gk189)* mutant at different stages of adulthood. (B) Food clearance assay in N2 (wt) and *glod-4 (gk189)* mutant after 72 hours of feeding. (C) Quantification of pharyngeal pumping (#/30 sec) in N2 (wt) after treatment, with either 150 µM of Arginine (control) or MG-H1. (D) Food clearance assay in N2 (wt) worms after treatment for 72 hours with either 150 µM of Arginine (control) or MG-H1. (E) Quantification of pharyngeal pumping with different concentrations of MG-H1. (F) Food racing assay in N2 (wt) and *glod-4 (gk189)* at different stages of adulthood towards OP50-1. (G) Food racing assay in N2 (wt) and *glod-4 (gk189)* towards OP50-1 mixed with MG-H1 vs OP50-1 mixed with MGO. Student t-test for A, B, C, E & F. One way ANOVA with Fisher’s LSD multiple comparison test for D & G. * p<0.05, ** p<0.01, *** p<0.005 and **** p<0.0001. Error bar ± SD.

### Tyramine regulates MG-H1 mediated feeding behavior via G-protein-coupled receptors (GPCRs) TYRA-2 and SER-2

Next, we sought to elucidate how MG-H1 increases the feeding behavior in worms. We performed an unbiased RNA sequencing approach to analyze the global transcriptome profile between control and *glod-4* knockdown worms (Suppl. File). Gene set enrichment analysis showed that the functional category of genes regulating feeding behavior was significantly upregulated in *glod-4* knockdown worms (> 2-fold enrichment score) (Figure 2A, red * marked GO category). This analysis supports our above observation that *glod-4* mutants have an altered feeding rate. Previous studies in *C. elegans* have documented the role of neurotransmitters in *C. elegans* feeding behavior [34] [35] [36] and we observed differential expression of 66 neurotransmitters and feeding genes (which comprises ∼19% of the total feeding and neurotransmission-related genes in *C. elegans*) (Figure 2B).

**Figure 2:**
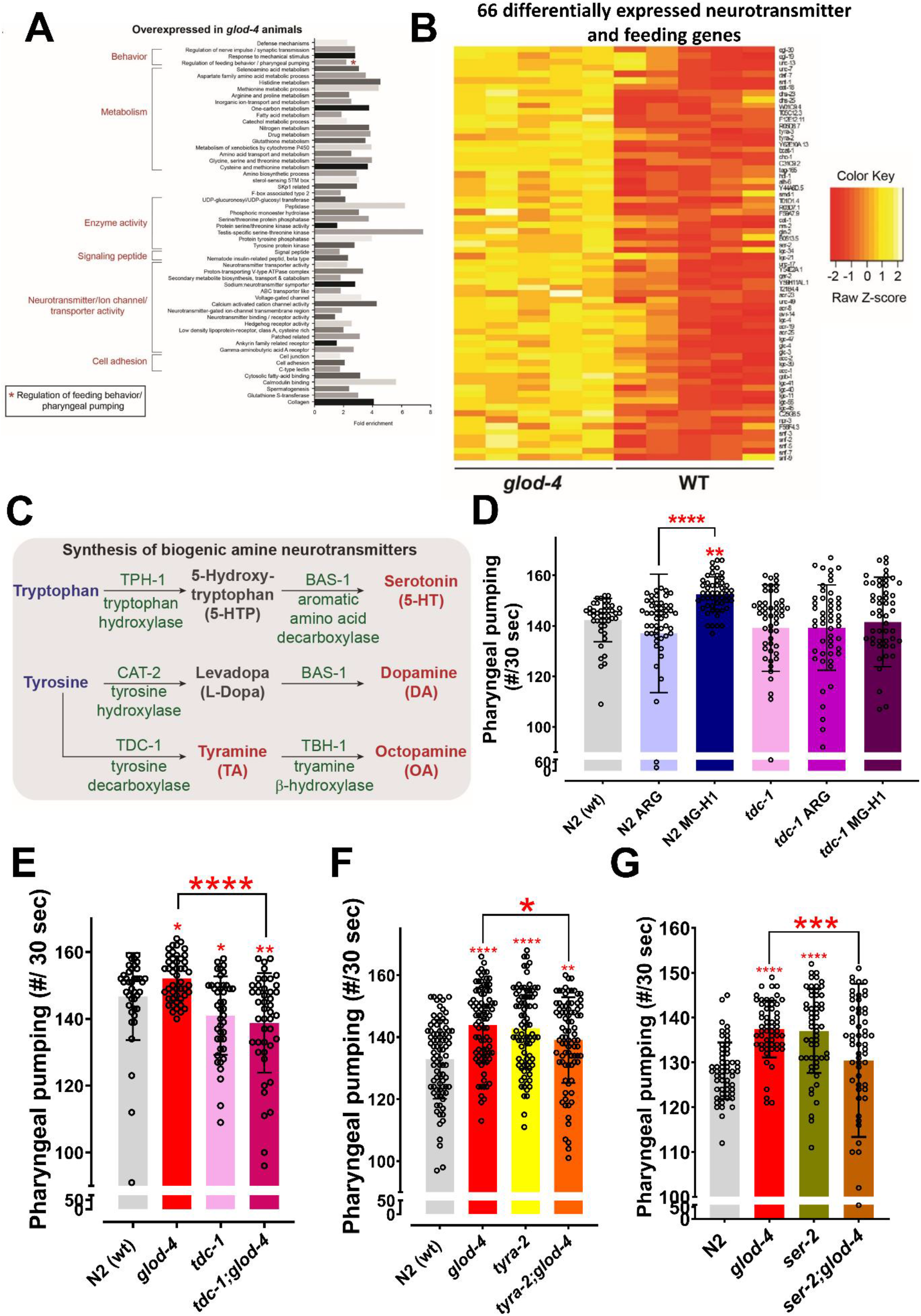
Role of *tdc-1* - tyramine receptors in mediating the MG-H1 induced feeding behavior. (A) Gene ontology analysis of differentially expressed genes in *glod-4* RNAi worms. (B) Differential expression of 66 neurotransmitters and feeding genes in *glod-4* RNAi background. (C) The flowchart shows the pathway of biogenic amine synthesis, which functions as a neurotransmitter. (D) Quantification of pharyngeal pumping in N2 (wt) and *tdc-1* mutant worms after 24 hrs of treatment of MG-H1. (E) Quantification of pharyngeal pumping in N2 (wt), *tdc-1, glod-4*, and *tdc-1;glod-4* double mutants. (F & G) Quantification of pharyngeal pumping in N2 (wt), *tyra-2, ser-2, tyra-2;glod-4* and *ser-2;glod-4* mutants. One-way ANOVA for D-G. * p<0.05, ** p<0.01, *** p<0.005 and **** p<0.0001. Error bar ± SD.

We next tested the involvement of these neurotransmitter genes in regulating MG-H1 mediated feeding behavior, and systematically analyzed MG-H1-induced feeding in the background of genetic mutants limited in producing different biogenic amines and neurotransmitters in *C. elegans* (Figure 2C + Figure Suppl. 2A). We found that mutation in *tdc-1*, the gene involved in synthesizing neurotransmitter tyramine, significantly suppressed the enhanced feeding phenotype in MG-H1 treated animals (Figure Suppl. 2A + Figure 2D). We also confirmed suppression of increased feeding rate in *tdc-1;glod-4* double mutant animals compared to *glod-4* (Figure 2E). Next, we checked putative receptors for tyramine that could potentially mediate downstream signaling. Receptors for tyramine and octopamine are well-studied G-protein-coupled receptors (GPCRs) [37] [38]. We screened seven GPCRs to identify the potential link in regulating tyramine-mediated increased feeding rate exhibited by *glod-4* mutant worms or MG-H1-treated worms. Observed results showed a mutation in *ser-2* and *tyra-2* suppresses enhanced feeding in MG-H1 treated animals (Figure Suppl. 2B). A similar reversal of feeding phenotype was observed in *tyra-2;glod-4*, and *ser-2;glod-4* double mutant strains (Figure 2F+2G). Our findings support the idea that MG-H1 induced overfeeding is mediated by tyramine signaling.

### GATA transcription factor *elt-3* acts upstream of *tdc-1 to* regulate MG-H1-mediated feeding behavior

To check for putative transcription factors (TFs) that could regulate the 66 differentially expressed genes in *glod-4* knockdown worms, we performed a motif-enrichment analysis (based on available ChIP-Seq data) (Figure 3A+3B). We chose the top five TFs (with a threshold of >18.75% target sequence match for TF-binding motif) for further screening. We knocked down each of the five TFs individually and checked for the rescue of the feeding behavior in worms exposed to 100 µM MG-H1 (Figure Suppl. 3A). Knocking down *pha-4* and *elt-3* suppressed the increase in pharyngeal pumping induced by MG-H1 treatment (Figure Suppl. 3A + Figure 3C). The *pha-4* gene is crucial for pharynx development, and loss of *pha-4* results in a morphological defect of the pharynx [39] [40]. Therefore, we chose to follow the results from *elt-3* knockdown worms with genetic mutants. Analysis of pharyngeal pumping in *elt-3;glod-4* double mutant showed that *elt-3* is essential to increase pharyngeal pumping observed in *glod-4* mutant worms (Figure 3D). To determine the role of *elt-3* in the tyramine signaling pathway, we performed a HOMER (Hypergeometric Optimization of Motif EnRichment) analysis and identified the binding site of *elt-3* on the *tdc-1* promoter, which suggested *elt-3* may potentially regulate *tdc-1* expression levels (Figure Suppl. 3B). This was further validated by reduced expression of *tdc-1* mRNA levels in the *elt-3* mutant worms (Figure 3E). Next, to check if the *elt-3* expression is changed on exposure to MG-H1, we treated wild-type N2 worms with MG-H1 and quantified mRNA levels of *elt-3*. We observed a moderate but significant increase in the *elt-3* expression (Figure 3F). Although *tdc-1* and *tyra-2* did not change significantly, expression levels of other tyramine receptors, *tyra-3* and *ser-2*, increased significantly after MG-H1 exposure (Figure 3F). Note that *ser-2* is necessary to mediate the increased pharyngeal pumping (Figure 2G). Together these experiments identified a key role for *elt-3* in tyramine-induced feeding increase in response to MG-H1.

**Figure 3:**
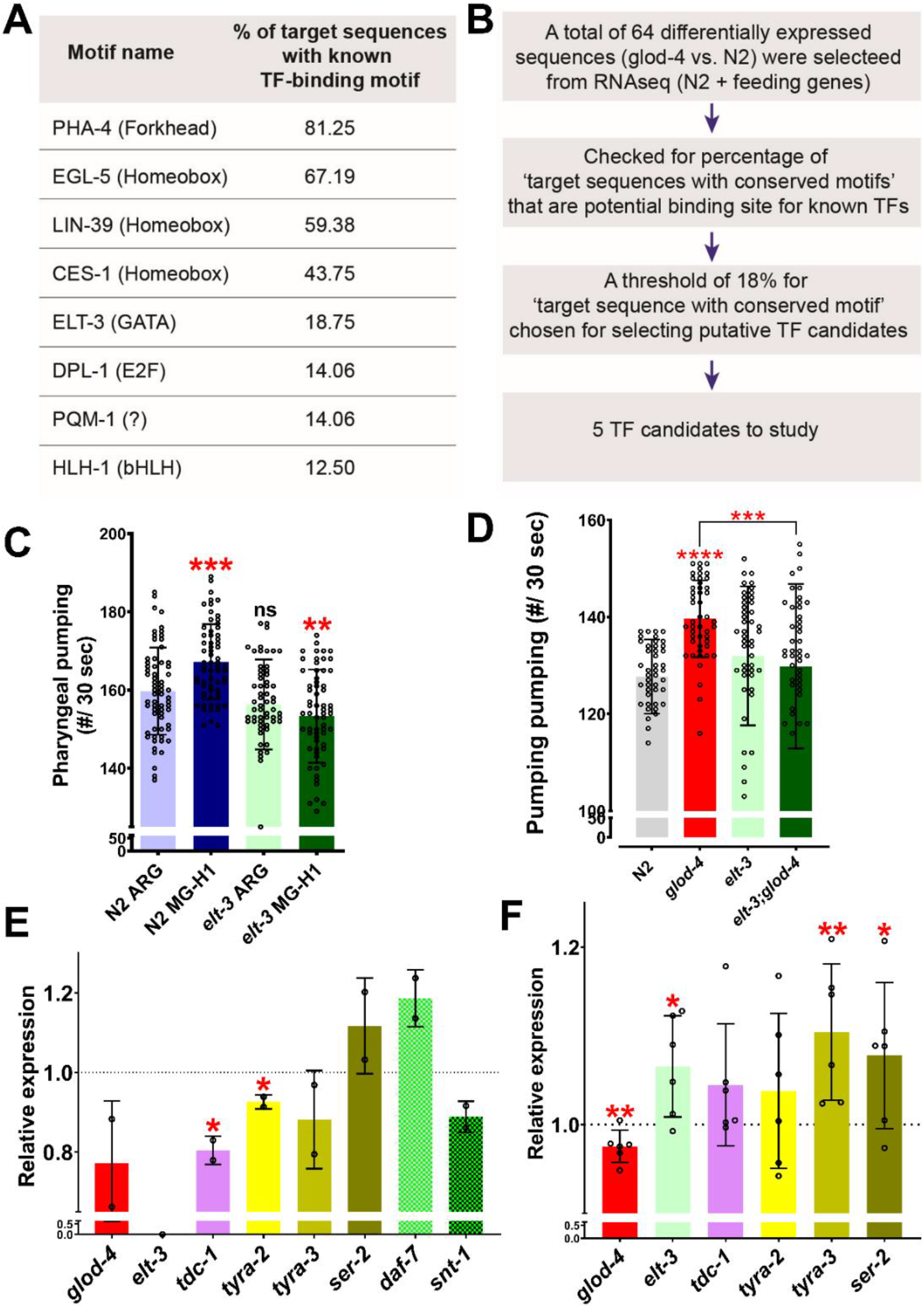
Role of *elt-3* transcription factor in regulating MG-H1 induced feeding in *C*. *elegans*. (A) List of transcription factors identified by motif analysis. (B) Flowchart demonstrating the method of identification of transcription factors. (C) Quantification of pharyngeal pumping after treatment with either Arginine or MG-H1 in *elt-3* mutants. (D) Quantification of pharyngeal pumping in N2 (wt), *glod-4, elt-3*, and double mutant worms. (E) Quantification of *elt-3* target genes in *elt-3* mutant worms. (F) Quantification of *elt-3* and tyramine pathway gene expressions in wild-type N2 (wt) worms after MG-H1 treatment. Horizontal dotted line indicate the normalized expression levels of genes in N2 (wt) and untreated control in E and F, respectively. One-way ANOVA for C+D. Student’s t-test for E+F. * p<0.05, ** p<0.01, *** p<0.005 and **** p<0.0001. Error bars ±SD.

### Absence of tyramine rescues α-DC mediated pathogenic phenotypes

Accumulation of α-DC in *glod-4* mutants results in pathogenic phenotypes including neurodegeneration and shortening of lifespan [17]. Here, chronic accumulation of MGO leads to the build-up of AGEs, thereby increasing feeding in *glod-4* worms (Figure 1). To test whether the pathogenic phenotypes exhibited by *glod-4* mutants result due to enhanced feeding, we compared the lifespan between wild-type N2 and *glod-4* worms in the genetic mutants that lack tyramine. The lifespan of *glod-4* was significantly increased upon inhibition of tyramine signaling in the *tdc-1;glod-4* double mutation (Figure 4A). In addition to rescuing lifespan and feeding rate, the lack of tyramine also resulted in the partial but significant rescue of neuronal damage in *glod-4* animals (Figure 4D+E).

**Figure 4:**
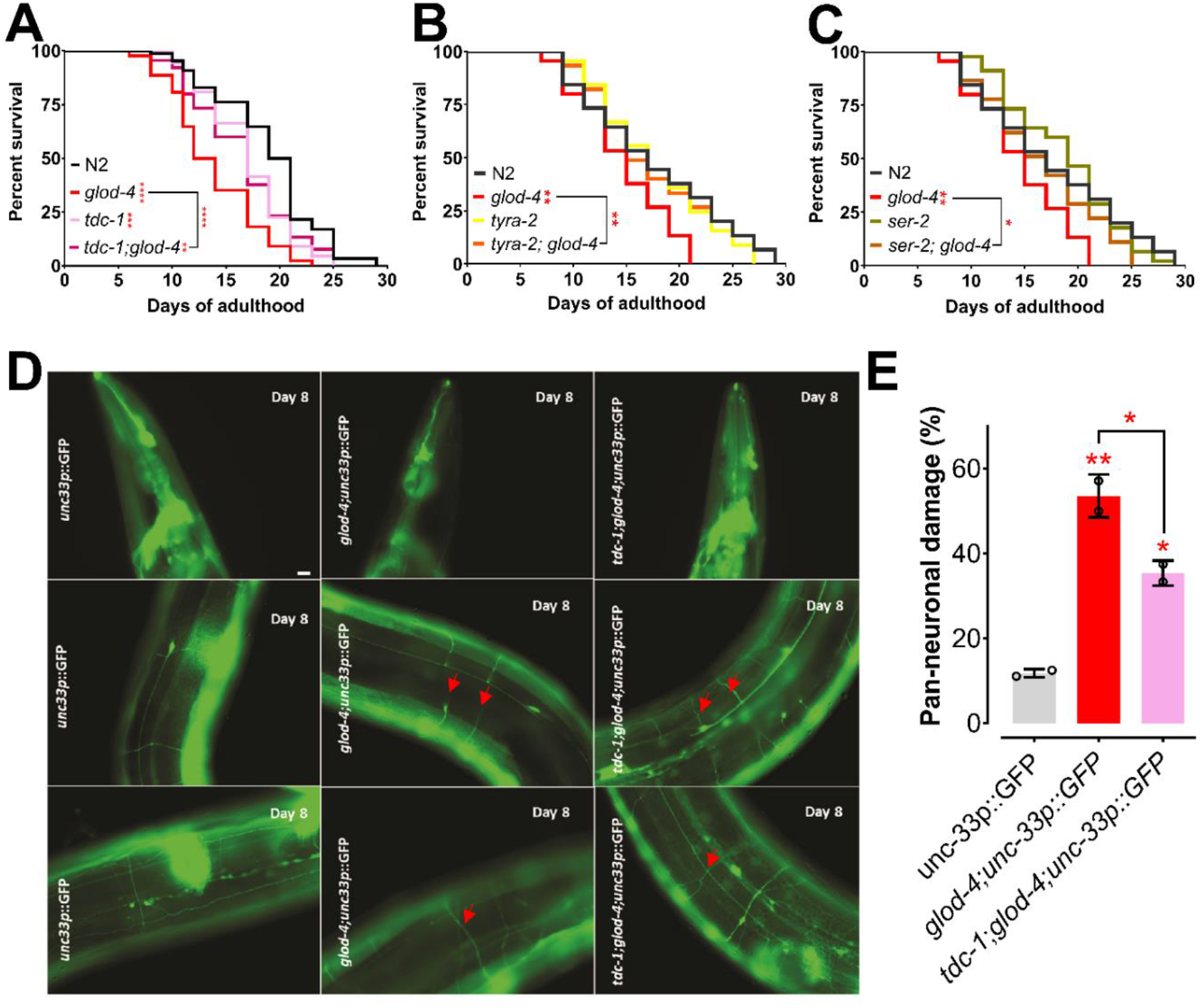
Suppression of *glod-4* phenotypes in *tdc-1;glod-4* double mutant. (A) Survival assay with N2 (wt), *tdc-1, glod-4*, and *tdc-1;glod-4* double mutants. (B) Survival assay with N2 (wt), *tyra-2, glod-4*, and *tyra-2;glod-4* double mutants. (C) Survival assay with N2 (wt), *ser-2, glod-4*, and *ser-2;glod-4* double mutants. (D) Image of worm neurons showing neuronal damage at day 8 of adulthood. (E) Quantification of neuronal damage with pan-neuronal GFP marker in *glod-4* vs. *tdc-1;glod-4* double mutants. Scale Bar – 10 µm. Log-rank (Mantel-Cox) test for survival assays. One-way ANOVA for (E). * p<0.05, ** p<0.01, *** p<0.005 and **** p<0.0001. Error bar ± SD.

Next, we tested if the absence of *tyra-2* and *ser-2* could also rescue the shortened lifespan of *glod-4* mutants. Lifespan increased significantly in the absence of either *tyra-2* or *ser-2* in double-mutant animals (Figure 4B+4C).

## DISCUSSION

Our observation that *glod-4* mutants run out of bacterial lawn faster than wild-type N2 animals during routine maintenance led to the elucidation of a novel signaling pathway that mediates AGEs-induced feeding behavior in *C. elegans*. Glyoxalases are enzymes involved in the detoxification of α-dicarbonyls, and we have previously characterized *glod-4* mutant, which lacks one of the glyoxalase enzymes, to accumulate increased levels of α-dicarbonyls and thereby AGEs [17]. In this study, using genetic mutants, synthesized AGEs, and functional genomics, we elucidate that AGEs (especially MG-H1) regulate feeding through tyramine signaling regulated by GATA transcription factor ELT-3. Also, MG-H1-induced hyper-feeding is independent of serotonin-mediated hyper-feeding [31] in *C. elegans* (Figure Suppl. 1B+1C). As both exogenous feeding of MG-H1 and the *glod-4* mutant, which enhances AGEs, led to increased feeding, we hypothesize that increased accumulation of AGEs is a potential stimulator of binge feeding. Thus, we studied changes in pumping rate by exogenous administration of MGO and AGEs. As previously reported by Ravichandran *et al*. 2018, MGO treatment did not change the pumping rate; however, MG-H1 and CEL increased the pumping rate in wild-type N2 worms (Figure Suppl. 1D) [41]. Further, a recent study demonstrated that treatment with sugar-derived AGE-modified Bovine Serum Albumin (BSA) accelerated the pharyngeal pumping rate[42]. In our study, we demonstrate that endogenous production and accumulation of AGEs via genetic mutation increase feeding and adversely affect lifespan.

Our detailed investigation of the time-dependent increase in pumping rate after MG-H1 treatment indicates a more robust and highly significant increase after 24 hours of treatment (Figure 1C). It is well established that AGEs are formed during cooking, browning the food during dry heating, making the food more appetizing [14]. Further, feeding is a multisensorial process regulated by several signaling pathways subjected to evolutionary adaptations [9]. Thus, we wanted to analyze the changes in sensory behavior of *C. elegans* induced by either endogenous accumulation of AGEs or by exogenous administration of MG-H1 with the food. Since the *glod-4* mutant lacks a glyoxalase system to detoxify methylglyoxal and leads to the accumulation of AGEs, the MG-H1 mediated signaling pathway can be responsible for the increased chemoattraction of *glod-4* mutant worms to food source OP50-1 (Figure 1F). It can be explained that including MG-H1 in bacterial lawn increased chemoattraction of wild-type N2 worms towards food, resulting in increased feeding of palatable MG-H1-mixed bacterial food OP50-1 (Figure 1G). However, unlike wild-type N2 worms, exogenous MG-H1 treatment had no further increase in the feeding rate or chemoattraction of *glod-4* mutant worms (Figure 1D+1G), indicating the maximum sensory modulation attained by the endogenous accumulation of MG-H1 in the *glod-4* mutant. Although our screening identified CEL, a lysine derived adduct of MGO, as another AGEs increasing the food intake, a detailed analysis is necessary to conclude the effect of CEL on feeding behavior (Figure Suppl. 1D).

We utilized RNA-seq data from *glod-4* knockdown worms to identify the novel signaling pathway that mediates AGEs-induced feeding in *C. elegans*. Since *glod-4* knockdown data are enriched with several genes regulating the synthesis of neurotransmitters and feeding (Figure 2A+2B), we performed suppression screening in mutant worms for genes involved in synthesizing biogenic amine neurotransmitters after MG-H1 treatment (Figure 2C and Figure Suppl. 2A+2B). Thus, our screen identified *tdc-1*, involved in the biosynthesis of tyramine, and tyramine receptors (*tyra-2* and *ser-2*) to mediate AGEs induced increased pharyngeal pumping (Figure 2D-2G) and Figure Suppl. 2). Tryptophan and tyrosine are the substrates for the synthesis of biogenic amines that have been implicated in modulating a wide array of behaviors in *C. elegans* [35, 43]. Tyrosine to Tyramine conversion in the presence of the enzyme tyrosine decarboxylase (TDC-1) followed by tyramine β-hydroxylase (TBH-1) is crucial for the synthesis of neurotransmitters tyramine and octopamine, respectively [43, 44]. Previous studies have shown the role of tyramine and its receptor (*ser-2*) in regulating feeding and foraging behavior in *C. elegans* [31] [45] [44] [46]. Further, the *tyra-2* receptor is expressed in MC and NSM pharyngeal neurons and discussed to potentially regulate pharyngeal pumping [31] [47]. Especially, tyramine has been shown to reduce pharyngeal pumping when applied exogenously to the worms [31]. Supporting previous findings [31], our observation shows increased pharyngeal pumping in *tyra-2* and *ser-2* single mutant worms (Figure 2F+2G); at the same time, *tdc-1* single mutants did not increase pumping (Figure 2E). Converse to our observation of *tyra-2* and *ser-2* single mutants, Greer *et al*. 2008 did not find any difference in the pumping rate of *tyra-2* and *ser-2* single mutants compared to wild-type N2 worms. However, the same study reported no changes in the pumping rate of *tdc-1* single mutant, similar to our results [44], which is also demonstrated by Li et al. 2012 [46]. Interestingly, double mutants of either *tdc-1* or its receptors (*tyra-2-*partial rescue and *ser-2*) with *glod-4* mutant significantly suppress the increased pharyngeal pumping observed in either *glod-4* or *tyra-2* or *ser-2* single mutants (Figure 2E-2G). It is to be noted that only two interneurons, namely RIM and RIC, uv1 cells near vulva and gonadal sheath cells [43] express the *tdc-1* gene, which is involved in the biosynthesis of tyramine; however, receptors of tyramine are expressed in distant tissues explaining an endocrine activity for tyramine neurotransmitter [31] leading to the multi-pathway mode of action to exert differential response which should be elucidated in future. Since *ser-3* mutant worms did not suppress the pumping (Figure Suppl. 2B) and *ser-3* has been demonstrated to be a receptor for octopamine [31], we conclude that octopamine is not responsible for mediating MG-H1 induced feeding in *C. elegans*.

Our suppressor screen for the upstream effector of the *tdc-1*-tyramine-*tyra-2/ser-2* pathway that mediates MG-H1-induced increased feeding identified the *elt-3* transcription factor (Figure 3C+3D). Thus, we examined whether *elt-3* TF regulates the *tdc-1, tyra-2*, or *ser-2*. Our analysis revealed that in *elt-3* mutant worms, *tdc-1* and *tyra-2* genes are significantly reduced (Figure 3E), concluding that *elt-3* TF regulates tyramine biosynthesis and receptor genes. In favor of the data, HOMER analysis identified *tdc-1* gene is potentially regulated by *elt-3* TF (Figure Suppl. 3B). Although *elt-3* TF is predominantly expressed in hypodermal cells, its expression is also reported in the pharyngeal-intestinal valve, intestine, few neurons (head neurons and mechanosensory PVD neuron), etc. (Wormbase.org). In accordance with *elt-3* expression in PVD neurons and head neurons, *tyra-2* is also expressed in PVD neuron [47] and *tdc-1* in RIM and RIC head interneurons, respectively, suggesting a possible direct/partial regulation of *tyra-2* and *tdc-1* expression by *elt-3*. Also, the *tyra-2* expression has been reported in pharyngeal MC neurons, which directly regulate pharyngeal pumping, [47] suggesting direct endocrine action of tyramine. Similarly, *ser-2* is expressed in pharyngeal muscle segment cells [45, 46, 48]. Though the *ser-2* expression is unchanged in *elt-3* mutant worms, *ser-2* expression is significantly increased in MG-H1 treated wild-type N2 worms (Figure 3F). Although the mechanism of MG-H1 induced expression of *ser-2* is unclear, it is evident that the *ser-2* genetic mutant can suppress the increased feeding in the *glod-4* mutant (double mutants) (Figure 2G), demonstrating an important role of the SER-2 receptor in mediating the MG-H1 induced feeding via tyramine. Further, *elt-3* expression levels significantly increased after MG-H1 treatment. Altogether our data strongly suggest the role of *elt-3-tdc-1*-tyramine-*tyra-2/ser-2* pathway in mediating enhanced feeding. Finally, it is essential to note that a few genes upregulated in the *glod-4* knockdown RNA-seq dataset have also been significantly upregulated after MG-H1 treatment, such as *ser-2* and *tyra-3*, validating that MG-H1 is a critical player in mediating adverse phenotypes observed in *glod-4* mutant worms.

Previously, we have demonstrated reduced lifespan, hyperesthesia, and accelerated neurodegeneration-like phenotypes observed in diabetic conditions, caused by excessive accumulation of α-dicarbonyls in *glod-4* mutant worms [17]. Lack of dicarbonyl detoxification by glyoxalases enzymes in *glod-4* mutant worms should result in the accumulation of AGEs, which at sufficient concentration act as signaling molecules to modulate the feeding behavior (Figure 1) by causing differential gene expression (Figure 2). The increased amount of dicarbonyl stress, thereby AGEs, is observed in several systemic diseases such as obesity, diabetes, cardiovascular and neurodegenerative diseases, among other age-associated diseases [1]. In diabetic patients, three times higher plasma levels of MGO have been reported, and is a leading cause of neuropathic pain [49] [50] [51]. The earlier reports in the literature show that the dicarbonyl levels correlate with diabetic complications. One of the major risk factors for diabetes is obesity [52], which is caused by overfeeding. Thus, exploring the regulatory pathways of feeding is essential to understand better and identify ways to modulate feeding behavior.

Here, we show that AGEs can modulate feeding behavior in evolutionary primitive model organisms, and it will be worth exploring this pathway in mammals. The transcription factor *elt-3* belongs to the GATA transcription factor family [53] Shobatake *et al*. 2018 report that GATA 2 and 3 transcription factors induce the expression of appetite regulator genes such as POMC and CART [54]. With the easy availability and unlimited access to modern-day processed food enriched in sugars and AGEs resulting in overeating, a significant cause of obesity pandemic, it is necessary to explore signals regulating feeding. Importantly, our study shows exogenous treatment with MG-H1 increases feeding in worms (Figure 1C+1D), indicating that a high AGEs diet in our day-to-day life can modulate feeding behavior in humans. It is well known that food cooked by grilling, broiling, roasting, searing, and frying accelerates the formation of AGEs in food; thus, methods are explored to cook food with fewer AGEs accumulation [55]. Further, increased caloric intake and changes in eating habits have been reported in a behavioral variant of Frontotemporal Dementia (FTP) [56] and medication of antipsychotic drugs [57].

Finally, we show that a lack of *tdc-1*-tyramine signaling rescues *glod-4* mutant phenotypes (lifespan and neuronal damage) (Figure 4). A strong association between worsening PD phenotypes with increased aggregation of α-synuclein and specific sites of increased glycation has been demonstrated in different genetic models with increased AGEs [58]. Thus, it will be interesting to investigate the role of the *tdc-1*-tyramine pathway in modulating pathways enhancing neurodegeneration and feeding. Recent research identified neurodegenerative diseases to be influenced by metabolism [59, 60] and *glod-4* mutants demonstrate increased neuronal damage, decreased lifespan, and increased feeding. Thus, it is essential to investigate the balance between energy metabolism to identify critical pathways to modulate the outcome of neurodegenerative diseases.

## MATERIALS AND METHODS

### Strains

Strains were either obtained from *Caenorhabditis* Genetic Center (CGC), Minneapolis, USA or National Bioresource Project, Tokyo, Japan and the following strains were used: N2 (wt), VC343 *glod-4(gk189)*, VC143 *elt-3(gk121)*, MT13113 *tdc-1(n3419), tyra-2(tm1846)*, RB1690 *ser-2(ok2103)*, MT9455 *tbh-1(n3247)*, CB1112 *cat-2(e1112)*, MT15434 *tph-1(mg280)*, DA1814 *ser-1(ok345)*, RB1631 *ser-3(ok2007)*, RB745 *ser-4(ok512)*, VC125 *tyra-3(ok325)*, and OH438 otls117[*unc-33p::gfp + unc-4(+)*]. Mutant strains are crossed to get the double mutants *elt-3;glod-4, tdc-1;glod-4, tyra-2;glod-4, ser-2;glod-4*, and *glod-4;unc-33p::gfp*. RNAi clones were obtained from Ahringer’s RNAi feeding library and the following were used: *pha-4, ces-1, elt-3, lin-39, egl-5*, and *tdc-1*.

### Growth and maintenance

Worms were cultured at 20°C for at least two generations on standard NGM agar plates seeded with 5X *Escherichia coli* OP50-1 bacterial strain (Broth culture of OP50-1 was cultured overnight at 37 °C at 220 rpm), which was propagated at RT for two days. For feeding RNAi bacteria, synchronized L1 larvae were transferred to NGM plates containing 3 mM of isopropyl β-D-1-thiogalactopyranoside/IPTG (referred to as RNAi plates) seeded with 20X concentrated HT115 bacteria (cultured overnight at 37 °C at 220 rpm), carrying the desired plasmid for RNAi of a specific gene or bacteria carrying empty vector pL4440 as control and allowed to grow on plates for 48 hrs. For drug assays, synchronized young adult worms (60 to 65 hrs from egg-laying) were transferred to NGM plates (with or without IPTG) with 20X HT115 RNAi bacteria or 5X OP50-1 bacteria (respectively), which are freshly overlayed by the desired drug (or vehicle control) that was air-dried and diffused. Final drug concentrations were calculated considering the total media volume on the NGM plates.

Note: For *glod-4* mutant animals, we found that the pathogenic phenotypes discussed in this paper are contingent on strictly maintaining an *ad-libitium* feeding regimen. Hence, care was taken not to allow the animals to starve by maintaining a low worm-to-bacteria ratio and transferring to fresh plates frequently (at least once every two days).

### Pharyngeal pumping assay

*C. elegans* pharyngeal pumping was measured using a Leica M165 FC stereomicroscope utilizing a modified previously established method [61]. Grinder movement in the terminal bulb was used as a read-out for the pumping rate phenotype. Pharyngeal pumping was recorded using a Leica M165 FC microscope; thus, obtained movies were played at X0.25 times the original speed and a manual counter was used to count the number of pumps for 30 sec. For quick pumping screening (pumping data in the supplemental figures), the pumping rate was counted in real-time for 30 seconds using a stopwatch and a manual counter focusing the grinder using an Olympus SZ61 stereomicroscope. 10 – 30 animals were counted per biological repeat and 2 to 3 repeats were obtained for each experiment and pumping data from all the repeats were combined for the presentation of data in the figures. At least one biological replicate was counted blind. Animals that did not pump during the recording time were eliminated from the analysis as well a few outliers were identified using the Gaussian distribution curve. Under exogenous drug treatment, animals were incubated in the drug at least 18-24 or until 48 hours before measurement of the pump rate. The drugs were overlaid on the NGM plate containing bacterial lawn and air dried before the addition of worms.

### Food clearance assay

Food clearance assay was performed following minor modification to established protocol by Wu et al. [62]. In brief, synchronized 20–25 L3-L4 stage worms were washed twice in S basal then once with S complete medium and transferred to a 96 well plate containing 160 µl assay medium (S-complete medium, growth-arrested OP50–1 at final OD 0.8 (at 600 nm), antibiotics, FuDR and either 150 µM Arginine or MG-H1 or 5 mM serotonin). Initial bacterial density was measured by obtaining OD at 600 nm. Following the indicated number of hours, bacterial density was measured at OD600 after a brief and gentle mixing using a multichannel pipet. For each experimental data point, at least six wells were measured (at least 120-150 worms in total), with the results shown being representative of at least two to three independent assays. The relative food intake was determined by the change in OD for each well, normalized to the number of worms. Under these conditions, ample OP50-1 was available for feeding throughout the analysis, and worms were maintained in the same wells for the entire duration of the experiment.

### Food race assay

The food race assay to evaluate *C. elegans* choice or attraction for a specific diet, a chemosensory behavior, was performed utilizing a previously established protocol [63]. For this assay, synchronized adult worms (50 per race) were spotted on a 60 mm NGM agar plate, freshly seeded with *E. coli* OP50-1 (with or without drug) approximately 2 cm from the edge of the Petri plate. Adult animals were aliquoted on the plate diametrically opposite to the food source to estimate the percentage of worms that reached the food source within 30 minutes. An illustration of the food-race assay has been provided (Figure Suppl. 1F).

### Organic synthesis of Advance Glycation End-products (AGEs)

#### Nδ-(5-hydro-5-methyl-4-imidazolon-2-yl)-ornithine (MG-H1) (3)

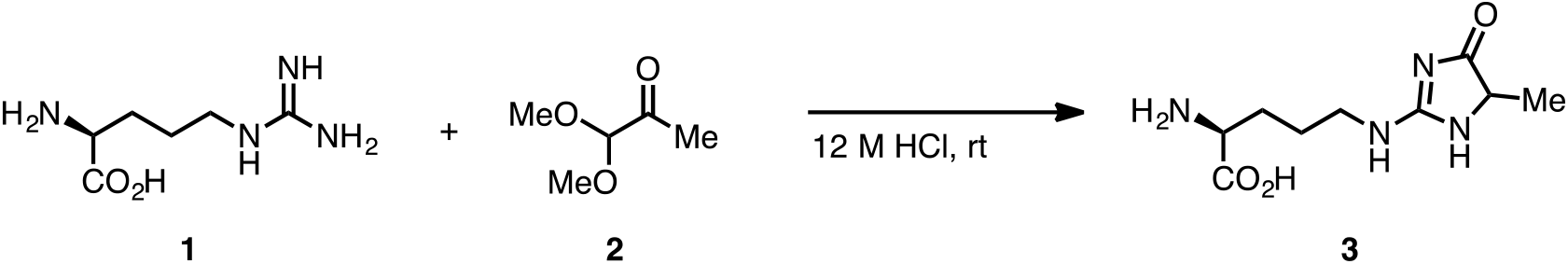

MG-H1 (**3**) was synthesized according to the literature procedure with a slight modification as follows; (*L*)-Arginine (**1**) (6.07 g, 34.8 mmol, 1 equiv) was dissolved in 12 M HCl (50 mL). To this was added methylglycol dimethyl acetal **2** (4.53 g, 38.3 mmol, 1.1 equiv). It was then stirred at room temperature for 11 hrs. At this time, the reaction mixture was diluted with water (200 mL) and concentrated *in vacuo*. The resulting dark-red solution was purified by SiO_2_-gel column chromatography (4:2:1 ethyl acetate:methanol:acetic acid) to give MG-H1 (**3**) as a yellow solid (5.23 g, 22.9 mmol, 66%). The spectroscopic data obtained are consistent with those previously reported in the literature ^[64]^.

#### Nε-carboxymethyl-lysine (CML) (7a) and Nε-(1-carboxyethyl)-lysine (CEL) (7b)

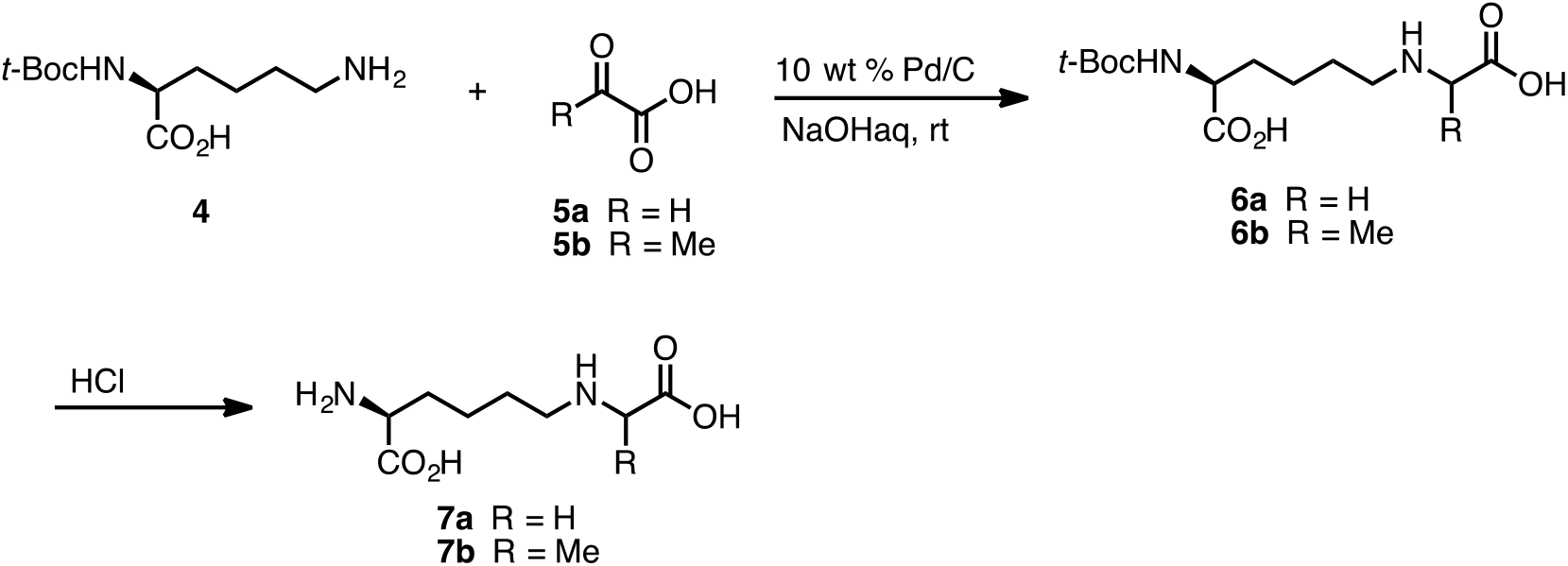

CML and CEL were synthesized according to the reported procedure ^[64]^ with a slight modification; To a 25 mL flask was added Nα-(*tert*-butoxycarbonyl)-*L*-lysine **4** (1.0 mmol, 1 equiv), palladium on carbon (10 wt.% loading, 100 mg, 0.94 mmol), and distilled H_2_O (7 mL). To this was added glyoxylic acid (120 mg, 1.3 mmol, 1.3 equiv) for CML synthesis or pyruvic acid (115 mg, 1.3 mmol, 1.3 equiv) for CEL synthesis. 1N NaOH_(aq)_ was added dropwise to make pH of this solution 9. A balloon filled with hydrogen gas was attached, and the resulting solution was stirred at room temperature for 14 hrs. At this time, the reaction mixture was filtered through a celite pad and the filtrate was concentrated *in vacuo*. Purification by SiO_2_-gel column chromatography (1:2 ethyl acetate:methanol) yielded **6a** (260 mg, 0.85 mmol) or **6b** (263 mg, 0.83 mmol), respectively. To this was added 1 N HCl_(aq)_ (3 mL), and it was then stirred at room temperature for 3 hrs. The resulting solution was concentrated *in vacuo* to give CML **7a** (164 mg, 0.80 mmol, 80%) or CEL **7b** (172 mg, 0.79 mmol, 79%). The spectroscopic data obtained are consistent with thosepreviously reported in the literature ^[64]^.

#### Synthesis of F-ly (12)

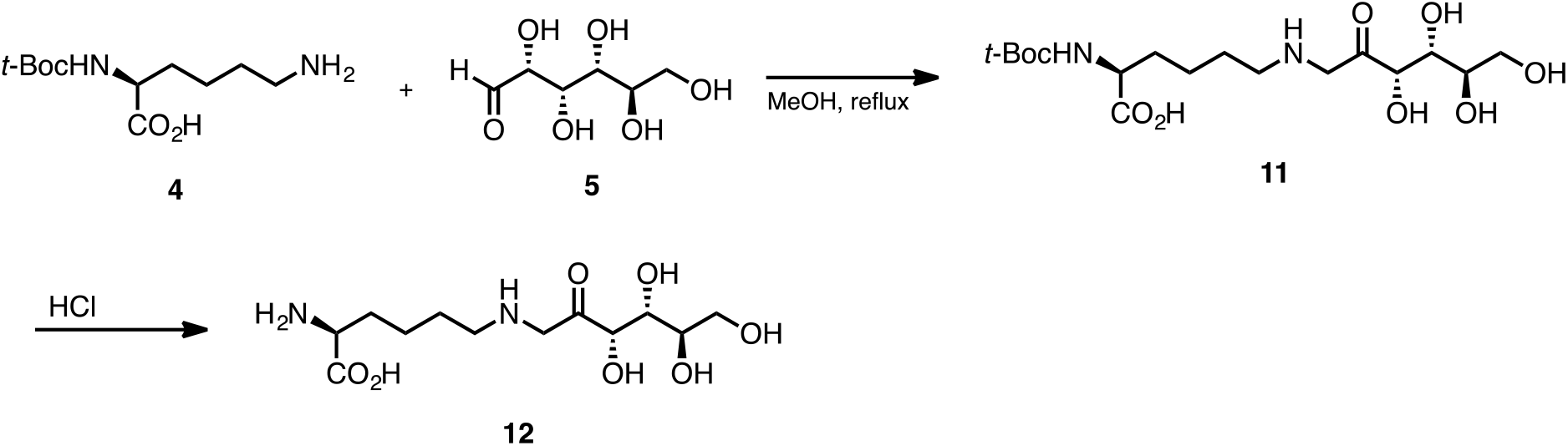

FLy (**12**) was synthesized according to the literature procedure with a slight modification as follows; To a 200 mL round-bottomed flask was added Nα-(*tert*-butoxycarbonyl)-*L*-lysine **4** (510 mg, 2.8 mmol, 1 equiv), D-(+)-glucose (6.15 g, 30.0 mmol, 10 equiv), and MeOH (90 mL). The condenser was attached, and it was refluxed for 7 hrs. After that, it was cooled to room temperature and concentrated *in vacuo*. The generated solid residue was purified by reversed-phase SiO_2_-gel chromatography (H_2_O only) to provide the desired compound **11** in 53% yield (599 mg, 1.47 mmol). This compound was reacted with 1 N HCl(aq) (3.5 mL) at room temperature and stirred overnight. After the concentration *in vacuo*, Fly **12** was obtained in 97% yield (438 mg, 1.42 mmol). The spectroscopic data obtained are consistent with those previously reported in the literature [65].

### Preparation of samples and methodology for RNAseq

RNA preparation for RNAseq was performed using the Qiagen RNeasy Mini kit (Cat. No. 73404). Total RNA extraction was performed from thirty-day 1 adult animals picked and collected in 20 µl M9 buffer per condition. 5 biological replicates were used for wild-type N2 and mutant animals. RNA-seq on the extracted total RNA was executed at the University of Minnesota Genomics Core (UMGC) using their sequencing protocol for HiSeq 2500 High Output (HO) mode and 50-bp Paired-end sequencing following Illumina Library Preparation. RNAseq coverage was ∼22 million reads per sample to perform downstream bioinformatics analyses.

### Bioinformatic analysis

RNAseq global transcriptome data were subjected to Gene Ontology (GO) based functional classification using the Database for Annotation, Visualization and Integrated Discovery (DAVID) v.6.8. We employed the heatmap2 (Galaxy Version 3.0.1) function from R ggplot2 package to visualize the bioinformatics data.

HOMER (Hypergeometric Optimization of Motif EnRichment) analysis was used to identify the transcription factors for the 66 differentially expressed gene from Figure 2B. A threshold of 18% was used to select potential transcription factors for further screening. Please refer to the flow chart in Figure 3B.

### Reverse transcription polymerase chain reaction (RT-PCR)

Total RNA was extracted from nearly 100 µl of tightly packed age-synchronized adult worm pellet collected in 1 ml TRIzol reagent provided by Qiagen RNeasy Mini kit (Cat. No. 73404) following manufacturer’s protocol. Subsequently, 1 μg total RNA was used as a template for cDNA synthesis. cDNA was synthesized using the iScriptTM cDNA synthesis kit (Bio-Rad, CA) following the manufacturer’s protocol. q-PCR was carried out using the PCR Biosystems Sygreen Blue Mix Separat -ROX (Cat. No. 17-507DB) in a LightCycler 480 Real-Time PCR system (Roche Diagnostics Corp., IN). Quantification was performed using the comparative ΔΔCt method and normalization for internal reference was done using either *act-5 or pmp-2*. All assays were performed with 3 technical replicates followed by 2-3 biological replicates. Following are qPCR primers used: 1) *act-5* gene primers are “TTCCAATCTATGAAGGATATGCCCTCCC and AAAGCTTCTCTTTGATGTCCCGGAC”. *pmp-2* primer pair is “ATCTTTCAAAGCCAATCCTCGAC and GAGATAAGTCAGCCCAACTCC”. *glod-4*: TGTTCTGAATATGAAAGTTCTTCGCCACG and GATGACGATTGCTCTATAATCATTACCCAACTC. *elt-3*: GCCGTTCAATATTTTTGAATTGAACCTTTCAAACTT and TTTTTTTCATCGGCTTCGGCTCG. *tdc-1*: CGACGAGTTGTTCCTGCTATT and CGGATGTTGCCAATGAGTTATTC. *tyra-2*: GGAAGAGGAGGAAGAAGATAGCGAAAGTAG and ATCTCGCTTTTCATCCGAGTCTTCATC. *tyra-3*: CATCGATGGCCGCTTGGTC and CTTGTTCTCGGGTATTTGAGCGGT. *ser-2*: GGAACAATTACGTACTTGGTAATTATTGCAATGAC and ATATCGCCACCGCCAGATCG. *snt-1*: CGGAAGCAGTAAAGCAAATAGCAACAAC and TCCCAGTTTGTTGCATAACCTTTTCTTTCA. And *daf-7*:AGAGTACCTTAAGAACGAAATTCTCGACCA and CCTTCTCCAGTAAGTCCCTATACATCTCC.

### Lifespan assay

Lifespan assays were performed in Thermo Scientific Precision incubators at 20°C. Timed egg laying was performed to obtain a synchronized animal population, which were either placed onto NGM plates seeded with 5X concentrated *E. coli* OP50-1. Post-L4 stage or young adult worms (60 to 65 hrs from egg laying) were added to FuDR (5-fluoro-2 deoxyuridine) NGM plates to inhibit the development and growth of progeny. After three days, animals were transferred to a new 60 mm NGM seeded with OP50-1 and scored every other day thereafter. 45-80 animals were considered for each lifespan experiment, and 2-3 biological replicates were performed. Animal viability was assessed visually and with gentle prodding on the head. Animals were censored in the event of internal hatching of the larvae, body rupture, or crawling of larvae from the plates [17].

### Assay for assessing neuronal damage

Neuronal damage was assayed using a pan-neuronal GFP reporter strain under different conditions on day 8 of adulthood. Animals were paralyzed using freshly prepared 5 mM levamisole in M9 buffer and mounted on 2% agar pads under glass coverslips. Neuronal damage was visually inspected under an upright Olympus BX51 compound microscope coupled with a Hamatsu Ocra ER digital camera. Images were acquired under the 40X objective. Neuronal deterioration was examined and characterized by loss of fluorescent intensity of nerve ring, abnormal branching of axon/dendrite, and thinning and fragmentation of axons and neuronal commissures [66]. Quantification and imaging of animals harboring damage were performed using the Image J™ software (http://imagej.nih.gov/ij/). To reduce experimental bias, this assay was performed genotype blind with 2 biological repeats.

### Statistical Analysis

All data analyses for lifespan, pharyngeal pumping assays, and gene expression were performed using GraphPad Prism (GraphPad Software, Inc., La Jolla, CA). Survival curves were plotted using the Kaplan-Meier method, and a comparison between the survival curves to measure significance (*P* values) was performed using Log-rank (Mantel-Cox) test. Two groups were compared for significance using an unpaired Student’s *t*-test. Multiple group comparison was performed by one-way ANOVA with either Fisher’s LSD or Dunnett’s multiple comparisons test, and Sidak’s multiple comparisons test was used to compare between specific groups. *P* values from the significance testing were designated as follows: \**P*<0.05, \*\**P*<0.005 and x002A;\*\**P*<0.0005.

## Data and material availability

All the data generated in this study are presented in the article. Synthetic AGEs can be obtained upon request.

## AUTHORS CONTRIBUTIONS

MMS, JC, and DS designed experiments, performed experiments, analyzed data, and wrote the manuscript. AKS, SG, MC, and BH performed experiments and analyzed data. RS and CR synthesized AGEs for the study. GL guided the study. PK conceived and guided the study, designed experiments, and obtained funding for the study.

## ACKNOWLEDGEMENTS

This work was supported by grants from NIH (R01AG061165 and R01AG068288) to PK and the Larry L. Hillblom Foundation (2021-A-007-FEL) to PK. We thank Professors Keith Blackwell, Suneil Koliwad, Malene Hansen and the members of the Kapahi lab for their valuable suggestions. We thank Dr. Feimei Zhu for her guidance in troubleshooting the food clearance assay. Dr. Kiyomi Kaneshiro for her guidance in organizing the RNAseq data. We also thank the Buck Institute’s morphology and imaging core for their valuable assistance.

## SUPPLEMENTAL FIGURES

**Figure 1 - Figure Suppl. 1:**
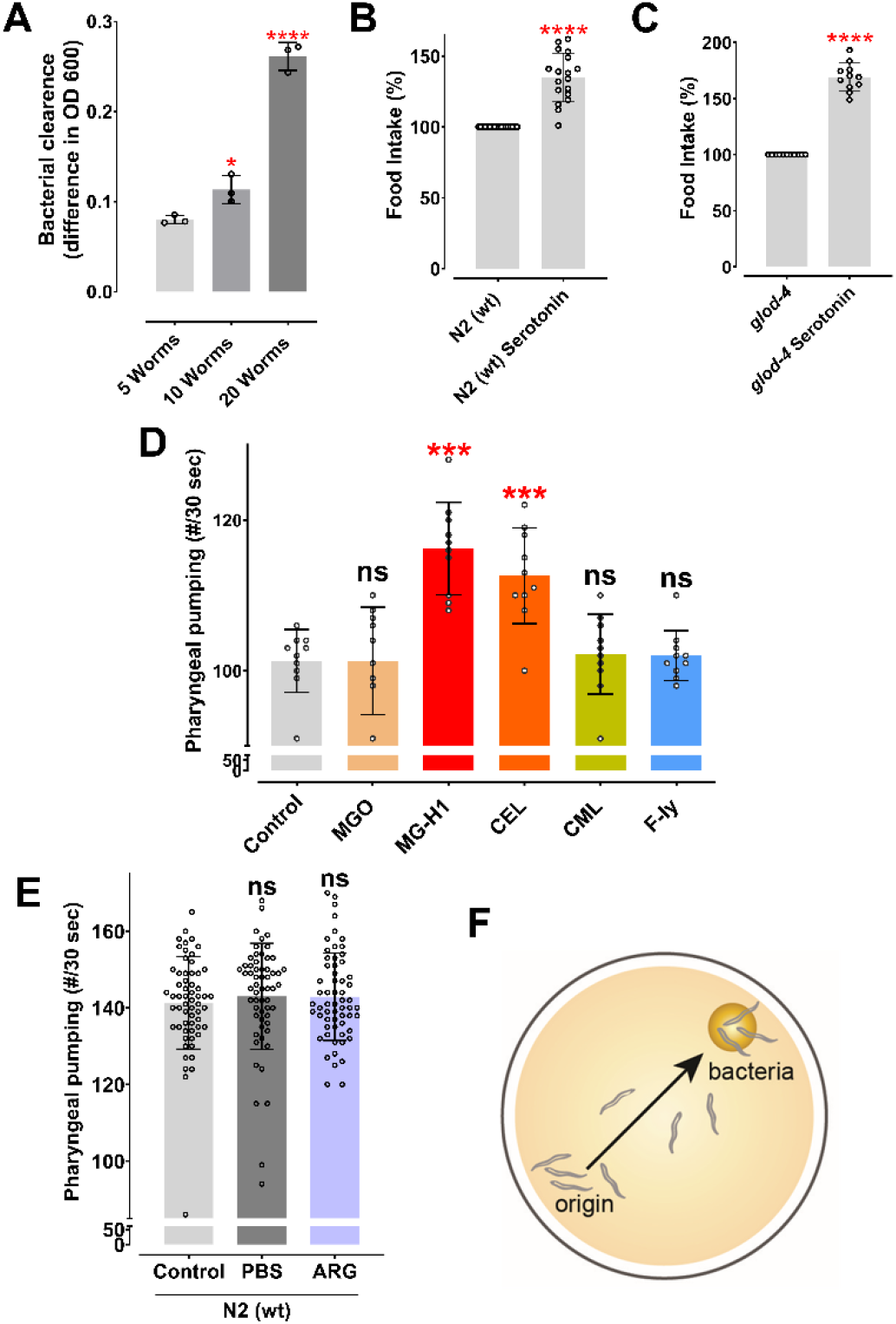
(A) Food clearance assay demonstrating increased food intake with an increasing number of worms. (B & C) Food clearance assay of wild-type N2 and *glod-4 (gk189)* mutant with 5 mM treatment of serotonin, respectively. (D) Quick visual quantification of pharyngeal pumping after treatment with different AGEs molecules at 100 µM for 12-18 hours. (E) Quantification of pharyngeal pumping in N2 (wt) worms after treatment with phosphate-buffered saline or 150 µM of Arginine for 24 hours. (F) Pictorial representation of food racing assay. Student’s t-test for B+C. One-way ANOVA for A+D. * p<0.05, ** p<0.01, *** p<0.005 and **** p<0.0001. Error bar ±SD.

**Figure 2 - Figure Suppl. 2:**
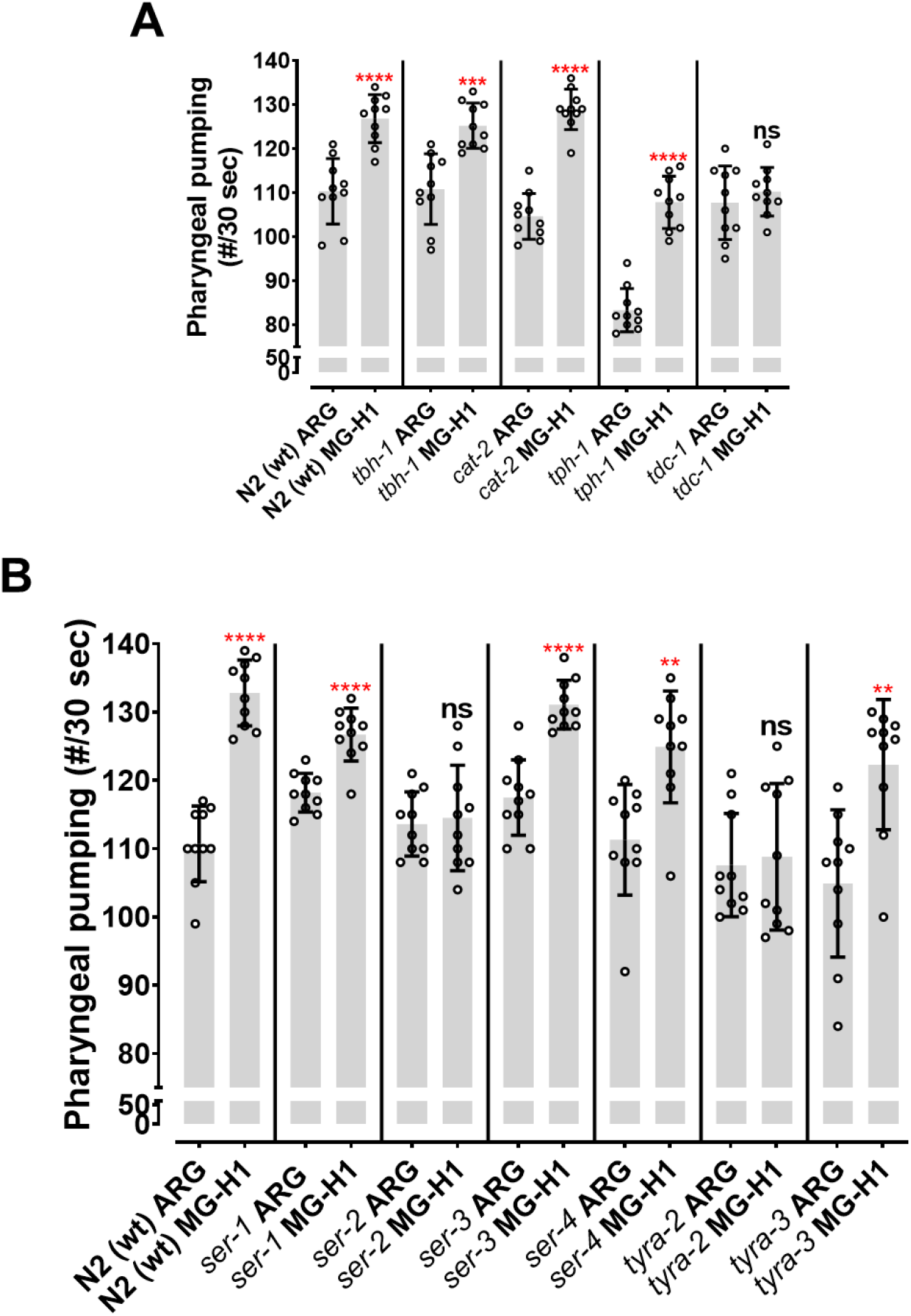
(A) Quantification of pharyngeal pumping (quick screening by visual counting) on mutants of enzymes involved in the biosynthesis of biogenic amines after MG-H1 treatment (suppressor screen). (B) Quantification of pharyngeal pumping (quick screening by visual counting) on receptor mutants involved in feeding behavior after MG-H1 treatment. Student’s t-test. * p<0.05, ** p<0.01, *** p<0.005 and **** p<0.0001. Error bar ±SD.

**Figure 3 - Figure Suppl. 3:**
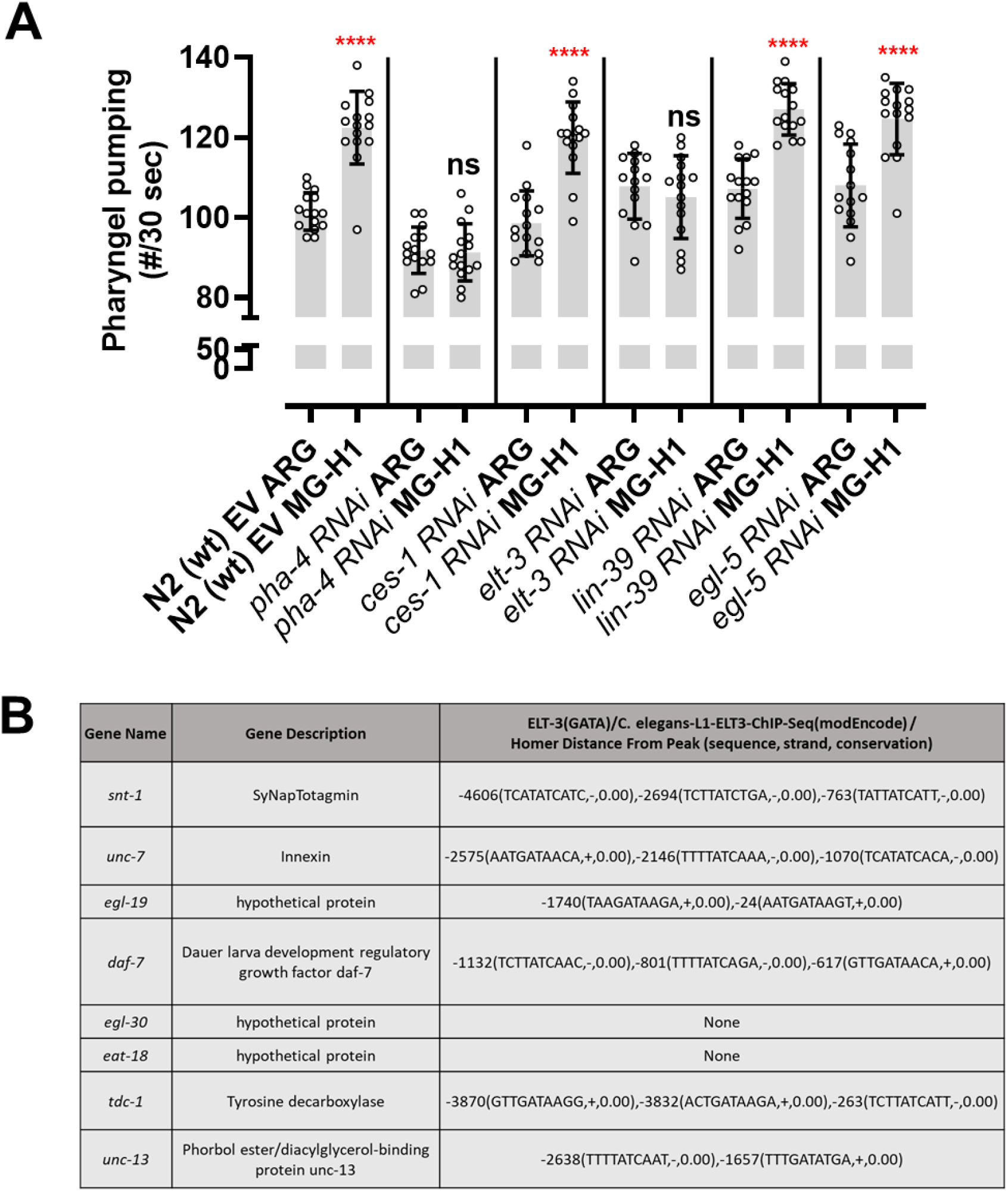
(A) Quantification of pharyngeal pumping (Quick visual counting), suppressor screen for top 5 transcription factors listed in Figure 3A. (B) List of genes obtained by HOMER analysis that are potentially regulated by the *elt-3* transcription factor. Student t-test in A. * p<0.05, ** p<0.01, *** p<0.005 and **** p<0.0001. Error bar ±SD.

